# Single species RNA purification with DNA nanoswitches

**DOI:** 10.1101/2020.07.07.191338

**Authors:** Lifeng Zhou, Cassandra Cavaliere, Andrew Hayden, Paromita Dey, Song Mao, Arun Richard Chandrasekaran, Jia Sheng, Bijan K. Dey, Ken Halvorsen

## Abstract

We report a novel method to purify individual RNA species using programmable DNA nanoswitches to facilitate RNA capture, release, and isolation. We validate sequence-based purification of microRNA, rRNA, and an mRNA fragment from total RNA, demonstrate multiplexing with two different RNA species, and show an application with downstream LC/MS analysis to identify RNA modifications. The simplicity, low cost, and low sample requirements of our method make it suitable for easy adoption.

---

Purification is a cornerstone of RNA research, arguably beginning in 1868 when Friedrich Miescher achieved the first nucleic acid (“nuclein”) purification^1^. Since then, many types of RNA with diverse functions have been discovered including messenger RNA (mRNA), catalytic ribozymes, self-splicing RNAs, and gene regulating RNAs^2^. The importance of RNA in human health is now well appreciated, especially with many viruses having RNA as their genetic information carrier. Additionally, recent discoveries of microRNAs, long noncoding RNAs (lncRNAs) and chemically modified RNAs have reshaped our understanding of the importance of RNA in biological processes and diseases^3,4^. The recent explosion of RNA research has made RNA purification increasingly important.

RNA purification is typically used to isolate all or most RNAs from a biological sample and is commonly done with organic extraction or spin columns. However, purification of specific RNAs is substantially more difficult, and magnetic beads-based purification is the only major approach for performing such isolations^5^. In this case, single-stranded DNA (ssDNA) capture probes on the beads target specific RNA sequences in cell lysates or total RNA samples. However, it is complex and expensive, with low yield and low specificity due to non-specific capture of RNA on the bead surface^6,7^.

Here we developed a novel detect-and-purify strategy for single species RNA purification (**Fig. 1a**) that overcomes many drawbacks of current approaches. Instead of capturing RNAs on a solid support (e.g. a magnetic bead), we use DNA nanoswitches that change their conformations upon binding the targeted sequence. Our nanoswitch is a linear double stranded DNA (dsDNA) with two ssDNA capture probes (oligo sequences presented in **Tables S1** to **S5**) that cause it to reconfigure to a looped dsDNA upon binding an RNA target (**Fig. 1b**)^8^. Our general strategy consists of three phases: 1) isolation of the RNA-looped nanoswitches by gel electrophoresis and gel excision, 2) digestion of DNA nanoswitches and gel pieces, and 3) removal of digested byproducts. First, the DNA nanoswitches capture target RNA molecules and become looped. Looped nanoswitches are separated and imaged using gel electrophoresis and isolated using gel excision. The nanoswitches are then digested using DNase I, which we found to work even on intact gel pieces (**Fig. S1**). The last step removes byproducts using a commercially available kit to dissolve the agarose gel pieces and then to purify the RNA from enzymes and nanoswitch fragments (**Fig. 1c**).

**Figure 1.**
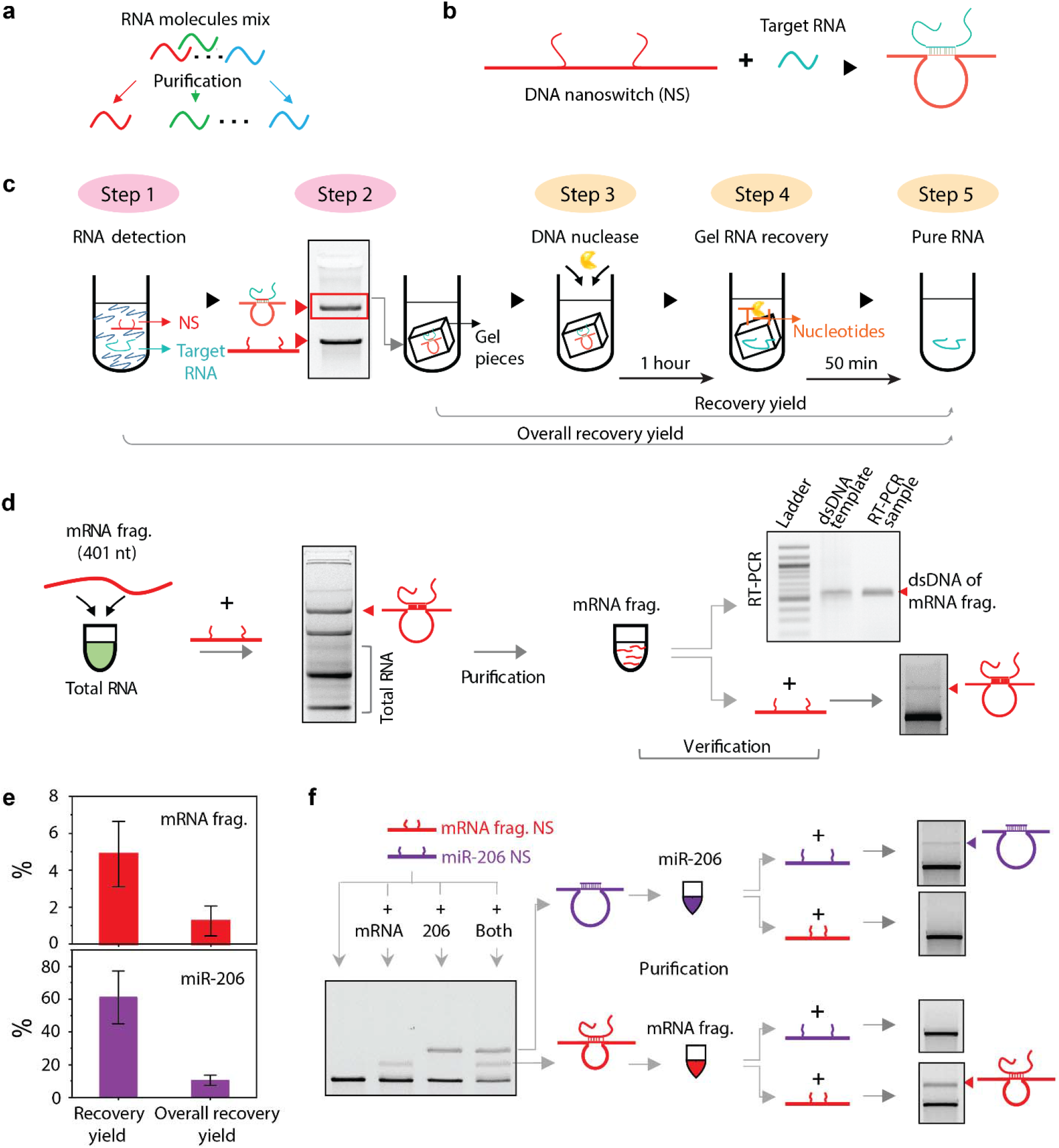
Single species RNA purification based on DNA nanoswitches. (**a**) Challenge of single species RNA purification. (**b**) The DNA nanoswitch converts from a linear to looped form in the presence of target RNA. (**c**) The purification workflow consists of detecting (steps 1-2) and purifying (steps 3-5) RNA. (d) Proof-of-concept validation for the purification of an mRNA fragment in total RNA with nanoswitch re-detection and verification by qRT-PCR. (**e**) The quantified purification yields of mRNA fragment and miR-206 based on qRT-PCR tests. (**f**) Multiplexed purification of mRNA fragment and miR-206 in single reaction. From left to right, scheme shows the multiplexed detection, purification and verification by redetection without cross contamination.

To demonstrate the purification, we chose two target RNA molecules with different lengths: a 401 nucleotide (nt) transcribed RNA fragment from the 3’ untranslated region of a *Drosophila* mRNA^9^ and a synthetic 22 nt microRNA-206 (miR-206). We first chose a target region (40 nt) on the mRNA fragment (**Table S1**). We designed four versions of nanoswitches with different capture probe lengths and found that the nanoswitch with 20 nt was the most efficient, with a detection limit of ~12.5 pM for the mRNA fragment (**Fig. S2**). For miR-206, we previously demonstrated sub-picomolar detection ability and adopted the same nanoswitch design here for the detection and purification^8^.

As proof-of-concept, we conducted the purification procedure for the mRNA fragment and miR-206 in water as well as spiked into total RNA to mimic a biological context and we successfully recovered the targets in both cases (**Fig. 1d and Fig. S3**). We demonstrated successful purification of the correct sequence by re-detecting the purified product with the nanoswitches (**Fig 1d**). To further validate the purification and quantify the yield, we performed qRT-PCR on the purified products. Both products were successfully amplified, and the mRNA fragment was consistent with the full-length DNA template used for transcription (**Fig. 1d and Fig. S4**). The final amounts determined by qRT-PCR were compared to the captured RNA and the starting RNA to determine the recovery yield and overall yield, respectively. We found overall yield of mRNA fragment and miR-206 to be 1.2% and 10.5% (**Fig. 1e, Fig. S5 and S6)**.

A unique feature of the DNA nanoswitches is that they can be programmed to enable multiplexing^8^, which we used here for simultaneous purification of multiple RNA molecules from a single reaction. This cannot be easily accomplished by other methods such as bead-based purification. To achieve this, we redesigned the location of the capture probes for the mRNA fragment nanoswitch to form a smaller loop (with faster migration) than the looped nanoswitch of miR-206 (**Fig. S7**). With this design, we showed detection and individual purification of the mRNA fragment and miR-206 at the same time (**Fig. 1f**). Redetection of the purified products with nanoswitches demonstrated the specificity of the purification method. Each nanoswitch only detected the correct purified target (mRNA or miR-206), indicating successful isolation of target RNA without notable cross-contamination (**Fig. S7**). Based on previous multiplexed detection, this should be scalable to at least 5 different RNA molecules in a single reaction^10^.

To demonstrate purification of RNAs from real biological samples, we developed multiplexed nanoswitches targeting the 5.8S and 5S ribosomal RNAs (rRNAs) from HeLa cell total RNA (**Fig. 2a**). The 5.8S and 5S subunits (156 and 121 nt respectively) are critical for protein translation^11^, and have been shown to contain chemical modifications including pseudouridine^12,13^. We chose target regions for each based on secondary structure analysis and designed nanoswitches to form a large loop for 5.8S rRNA and a small loop for 5S rRNA (**Fig. S8**). We showed clear multiplexed detection of the 5.8S and 5S rRNAs directly from 250 ng total RNA from HeLa cells (**Fig. 2b**). Following our established workflow, we separately purified each rRNA species from 2 μg total RNA of HeLa cell (**Fig. S8**) and confirmed successful purification of the correct rRNA in each instance (**Fig. 2c**).

**Figure 2.**
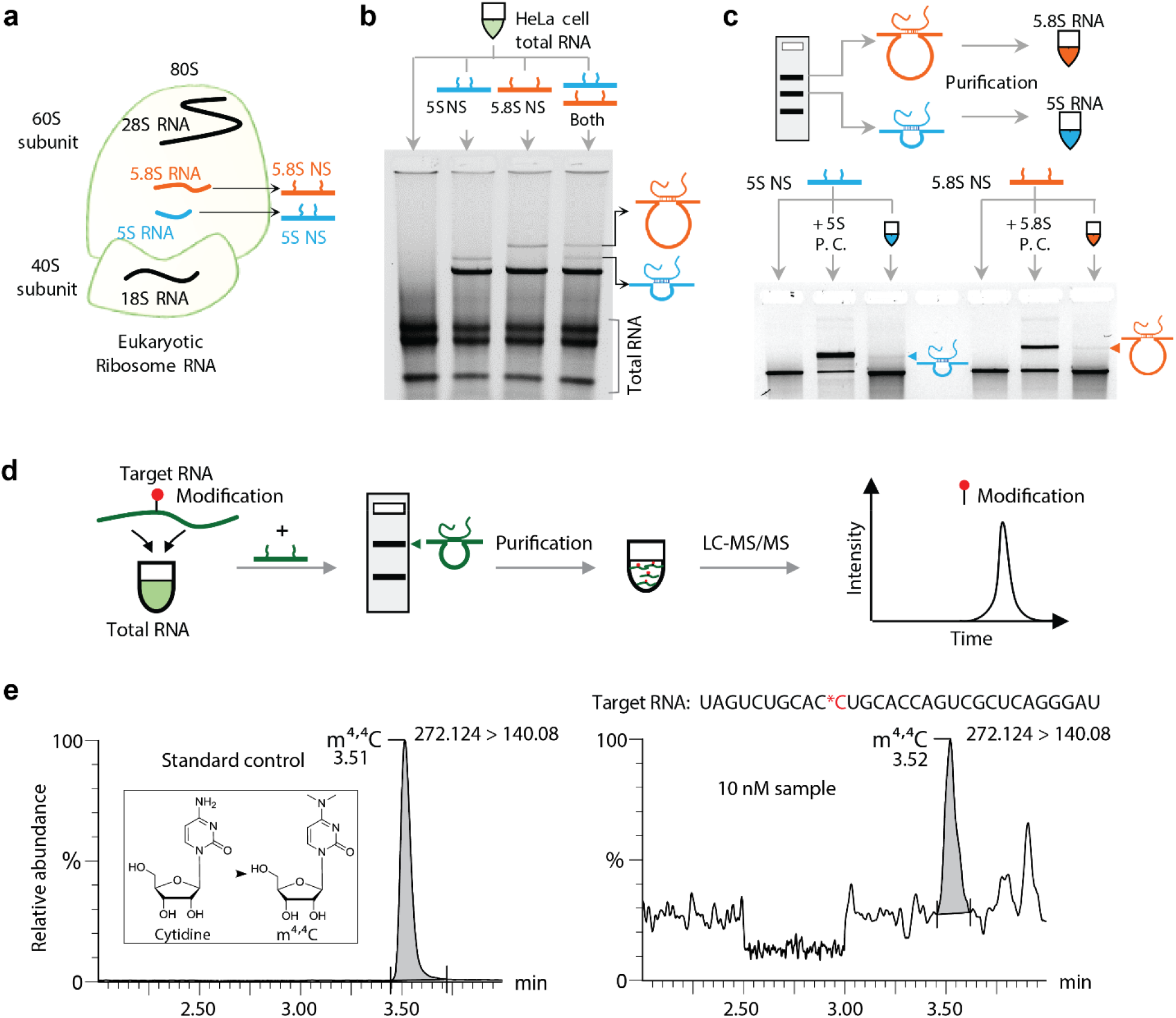
Biological applications of single RNA species purification. (**a**) Components of eukaryotic ribosome. (**b**) Multiplexed detection shows separate detection of 5.8S and 5S rRNAs as different loop sizes. (**c**) Multiplexed purification demonstrates individual isolation of the 5.8S and 5S rRNA molecules without observable cross-contamination. (**d**) The process of purification and verification by LC-MS/MS of modified RNAs. (**e**) Modification probing of the purified RNA molecules by LC-MS/MS. Left: m_2_^4^C modification standard and right: purified RNA from 10 nM sample. The target RNA molecule has a m_2_^4^C modification on a cytosine (sequence shown above graph on right, * indicates position of modification).

One compelling application of our approach is to study how the >100 natural RNA modifications can alter biological functions of particular RNAs^4^. For example, N6-methyladenosine (m^6^A) affects the stability of mRNA and protein translation^14^, rRNA modifications can influence translation efficiency^15^, and some RNA modifications appear in response to viral infection^16^. One of the gold standard methods for measuring RNA modifications is ultra high-performance liquid chromatography-tandem Mass Spectrometry (UHPLC-MS/MS)^17^, but this method typically uses digested RNA and loses sequence information. Used in conjunction with our purification method, RNA modifications could be measured on specific RNA sequences (**Fig. 2d**). To demonstrate this application, we chose the m^4,4^C (*N*^4^, *N*^4^-dimethylcytidine) modification (**Fig. 2e**), which has relevance in viral infection and to our knowledge can only be identified by mass spectrometry methods. We synthesized and a PAGE purified short (31 nt) RNA with a single m^4,4^C modification, and screened two nanoswitch designs since the modification can influence base pairing (**Fig. S9**). To mimic biological samples, we spiked the modified RNA at different concentrations (10 nM and 1 nM) into 2 μg of total RNA from HeLa cells and performed the purification (**Fig. S9**). We showed detection of the modification from the purified samples at both concentrations, demonstrating that our method can be applied to extract target RNA with chemical modifications with enough material for downstream UHPLC-MS/MS testing (**Fig. 2e and S10)**.

Our novel “sense and purify” method for single species RNA purification enables robust purification of diverse types of RNA from microgram scale samples in just a few hours. The capture probes of our nanoswitches can be readily programmed for multiple purification targets without significant effort. Our approach avoids the downsides of surface binding approaches present in bead-based methods, provides visual feedback of the process for troubleshooting, and can be performed at low cost on the benchtop (**Table S6**). We expect that future improvements could increase the overall yield and purification scale, expand multiplexing, and further reduce the cost and time for processing. Some other techniques could potentially speed or improve the collection of looped nanoswitches such as the BluePippin gel cassette^18^ or better spin columns (DNA gel extraction columns suffered from low yield - **Fig. S11**).

Analysis of chemical modifications of RNA is a particularly attractive application. Mass spectrometry techniques can detect modifications at below fmol scale but tend to lose sequence information. By purifying single RNA species for MS^17^, cryo-EM analysis^19^ or epitranscriptome sequencing technologies^20^, it will be easier to determine which particular RNAs contain modifications. It can be seen from the history of scientific literature that advances in purification tend to precede new discoveries (e.g. Dr. Meischer’s isolation of DNA in 1868). We hope that our approach will similarly facilitate new discoveries of RNA science.

## Materials and Methods

### DNA nanoswitches

All DNA nanoswitches were designed and fabricated according to the protocol presented in reference^8^. Briefly, circular M13 ssDNA (New England Biolabs) was linearized by enzyme BtsCI (New England Biolabs). The backbone and detection oligos (IDT DNA) were mixed with the linearized M13 with about 10× excess (all oligos used in this research are presented in **Table S1 to S5**). All nanoswitches were self-assembled in a thermal cycler with a thermal annealing protocol (90 to 25 °C, 1 °C/min). After fabrication, DNA nanoswitches were purified by either HPLC^8^ or by PEG precipitation^8^ as described previously. Purified nanoswitches were resuspended in 1× PBS and their concentrations were measured using a Nanodrop 2000.

### RNA samples

The 401 nt mRNA fragment was produced by *in vitro* transcription (NEB, HiScribe™ T7 High Yield RNA Synthesis Kit) from a DNA template (**Table S1 and S2**), which was a gift from Prof. Prashanth Rangan. The microRNA target (miR-206) was commercially synthesized (IDT DNA). Total RNA from HeLa cells was purchased from BioChain Institute Inc.

RNA strand containing the m^4,4^C modification was synthesized at 1.0 mmol scale by solid phase synthesis using an Oligo-800 synthesizer. After synthesis, the oligos were cleaved from the solid support and fully deprotected with AMA (ammonium hydroxide:methylamine solution = 1:1) at 65 °C for 45 min. The amines were removed by Speed-Vac concentrator followed by Triethylamine trihydrofluoride (Et3N.3HF) treatment for 2.5 h at 65 °C to remove the TBDMS protecting groups. Cooled down to room temperature the RNA was precipitated by adding 0.025 mL of 3 M sodium acetate and 1 mL of ethanol. The solution was cooled to −80 °C for 1 h before the RNA was recovered by centrifugation and finally dried under vacuum. The RNA strands were then purified by 15% denaturing polyacrylamide gel electrophoresis (PAGE) and were desalted, concentrated and lyophilized before redissolving in RNase free water.

### Nanoswitch detection assays

Unless otherwise noted, all nanoswitch detection and assays were performed by incubating the nanoswitches with target RNA in 1× PBS and 10 mM MgCl_2_ in a thermal annealing ramp (40 to 25 °C, 0.1 °C/min) for about 12 hours. Nanoswitch concentrations varied from ~0.1-1 nM as noted in figure captions, with typically high nanoswitch concentrations for high capture in the purification process and low concentrations for validation assays. Validation assays also included 200 nM of “blocking oligos” to minimize RNA sticking to the tubes. GelRed DNA stain and a Ficoll-based loading dye were added to the final samples at a 3.3× and 1× final concentrations, respectively. Gel electrophoresis was performed using 0.8% agarose gels in 0.5 × TBE buffer. Gels were run at 4°C at 60-75V for 45 minutes to 2 hours depending on the assay.

### RNA purification protocol

After performing the nanoswitch detection assay and running gel electrophoresis, the looped nanoswitch gel bands were excised on the image platform of a Gel Doc XR+ System (Bio-Rad) by using disposable plastic gel cutting tool and razor (Sigma-Aldrich). All gel bands were diced into small pieces before transfer into 1.5 ml tubes. Then, the gel pieces were submerged in 1× DNase I buffer (NEB) and 4 U DNase I (NEB) was used for each 200 μl to digest the DNA nanoswitches at 37 °C for 1 hour. Following DNase I digestion, Zymoclean Gel RNA recovery kit (Zymo research) was used to recover and clean the target RNA. The manufacturer’s instructions were followed, except for doing two sequential elutions for the last step in 10 μl nuclease-free water each. Totally, for each column, we obtained 20 μl RNA sample. An optional 15-minute heating step at 90 °C was used to destroy any residual DNase I before downstream re-detection by the DNA nanoswitches.

### Modification analysis by LC-MS

Measurement of the level of m^4,4^C was performed by ultra-performance liquid chromatography coupled with tandem mass spectrometry (UHPLC-MS/MS) using a method similar to that previously described in reference^21^. After the purification of the RNA with m^4,4^C modification, we first dried the sample in a universal vacuum system (Savant UVS 400) and then resuspended in RNase-free water and digested the RNA into nucleotides by Nucleoside Digestion Mix (NEB, M0649S) in 15 μl volume and used 10 μl for the test. The UHPLC-MS/MS analysis was accomplished on a Waters XEVO TQ-S™ (Waters Corporation, USA) triple quadruple tandem mass spectrometer equipped with an electrospray source (ESI) maintained at 150 °C and a capillary voltage of 1 kV. Nitrogen was used as the nebulizer gas which was maintained at 7 bars pressure, flow rate of 1000 l/h and at a temperature of 500°C. UHPLC-MS/MS analysis was performed in ESI positive-ion mode using multiple-reaction monitoring (MRM) from ion transitions previously determined for m^4,4^C (m/z 272 > 140). A Waters ACQUITY UPLC™ HSS T3 guard column (2.1 x 5 mm, 1.8 μm) attached to a HSS T3 column (2.1 x 50 mm, 1.7 μm) was used for the separation. Mobile phases included RNAse-free water (18 MΩcm^-1^) containing 0.01% formic acid (Buffer A) and 50% acetonitrile (v/v) in Buffer A (Buffer B). The digested nucleotides were eluted at a flow rate of 0.4 ml/min with a gradient as follows: 0-2 min, 0-10% B; 2-3 min, 10-15% B; 3-4 min, 15-100% B; 4-4.5 min, 100 %B. The total run time was 7 min. The column oven temperature was kept at 35 °C and sample injection volume was 10 μl. Three injections were performed for each sample. Data acquisition and analysis were performed with MassLynx V4.1 and TargetLynx. Calibration curves were plotted using linear regression with a weight factor of 1/concentration (1/x).

### qRT-PCR assays

cDNA synthesis of purified mRNA fragment was carried out using the iScript cDNA Synthesis Kit (Bio-Rad) as instructed. The various amount of known mRNA fragments (0.0625 nM, 0.125 nM, 0.25 nM, 0.5 nM, and 1 nM) were similarly converted into cDNAs and used to generate a standard curve. Then, qRT-PCR was carried out using SYBR green PCR master mix (Bio-Rad) in a Bio-Rad Realtime Thermal Cycler using purified fragment specific primers (Forward-TGTTTGCTTTCGTGAAAACTCGCAT; Reverse-ACATTAGGTGCAATACCGAAGGCA). The concentration of unknown purified fragments was derived from the known amount of fragments used to generate standard curves.

cDNA synthesis of purified miR-206 was carried out using miRCURY LNA RT Kit (Qiagen) as described. Briefly, every 10 μl RT reaction contained 2 μl of 5× reaction buffer, 4.5 μl of RNase free water, 1 μl of 10× RT enzyme mix, 0.5 μl of RNA spike in SP6 and 2 μl of template microRNA. The reactions were thoroughly mixed in 0.2 ml tubes and subjected to cycling: 60 min at 42 °C, 5 min at 95 °C. Similarly known amount of synthetic miR-206 (0.0625 nM, 0.125 nM, 0.25 nM, 0.5 nM, and 1 nM) were converted into cDNAs and used to generate a standard curve for calculating unknown miR-206 concentration. qRT-PCR for microRNA was done following the instructions of miRCURY SYBR Green PCR Kit (Qiagen). Briefly, for every 10 μl reaction, 5 μl of 2× miRCURY SYBR Green Master Mix, 2 μl of Nuclease Free Water, 1 μl of miR-206 PCR Primer Mix, and 2 μl of cDNA Template were used. The reactions were mixed well and dispensed into PCR plate (10 μl/well), which was then placed in a Bio-Rad Realtime Thermal Cycler using the cycling protocol: after 2 minutes at 95 °C, 40 repetitions of 10 seconds at 95 °C and 60 seconds at 56 °C,. The melt curve analysis was done at 60-95 °C. The concentration of unknown purified miR-206 was derived from the known amount of miR-206 used to generate standard curves.

## Acknowledgements

We thank Prash Rangan for material and assistance with the mRNA fragment, Simon Chi-Chin Shiu for performing a validation experiment and providing manuscript suggestions, and Dr. Qishan Lin for LC-MS/MS tests and Jibin Abraham Punnoose for manuscript suggestions.

## Funding

Research reported in this publication was supported by the NIH under award R35GM124720 to K. H. The content is solely the responsibility of the authors and does not necessarily represent the official views of the NIH.

## Author contributions

L. Z. and K.H. designed the major experiments. L.Z., C.C., and A. H., performed nanoswitch experiments. P. D. and B.K.D. designed and performed qRT-PCR experiments. S. M. and J. S. designed and synthesized the modified oligonucleotide. L.Z. wrote the first draft of the manuscript. L.Z., A.R.C and K.H. co-wrote later drafts of the manuscript with comments from other authors.

## Competing finical interests

K.H. and A.R.C. have intellectual property related to DNA nanoswitches. All other authors declare that they have no competing interests.

## Data and materials availability

All data needed to evaluate the conclusions in the paper are present in the paper and/or in the Supplementary Materials. Additional data related to this paper may be requested from the authors.

## Notes

### Competing Interest Statement

LZ, ARC, and KH have intellectual property related to DNA nanoswitches.  All other authors declare no competing interests.

