## Supplemental Information for "Single species RNA purification with DNA nanoswitches"

***Contents***

**Figure S1.** Digestion of DNA nanoswitches in gel pieces.

**Figure S2.** Optimization of the detection arm length and sensitivity test of DNA nanoswitch of mRNA fragment.

**Figure S3.** Proof-of-concept single species RNA purification.

**Figure S4.** Length validation for mRNA fragment.

**Figure S5.** Quantification of the purification yield of mRNA fragment based on qRT-PCR test.

**Figure S6.** Quantification of the purification yield of miR-206 based on qRT-PCR test.

**Figure S7.** Gel images of the multiplexing purification of mRNA fragment and miR-206.

**Figure S8.** Detection and purification of 5.8S and 5S rRNA.

**Figure S9.** Purification of RNA with chemical modification.

**Figure S10.** Purification of RNA molecules with chemical modification.

**Figure S11.** Test of DNA gel extraction spin columns.

**Table S1.** Sequence of target mRNA fragment and oligos used for the design of nanoswitches.

**Table S2.** Sequence of target miR-206 and oligos used for the design of nanoswitches.

**Table S3.** Sequences of target ribosomal RNA 5.8S and 5S and oligos used for the design of nanoswitches.

**Table S4.** Sequence of synthesized RNA with m^4,4^C chemical modification and oligos used for the design of nanoswitches.

**Table S5.** Backbone used for the design of nanoswitches and blocking oligo used in some of the detection assays^1,2^.

**Table S6.** Cost of the DNA nanoswitch assay^1^.

**References**

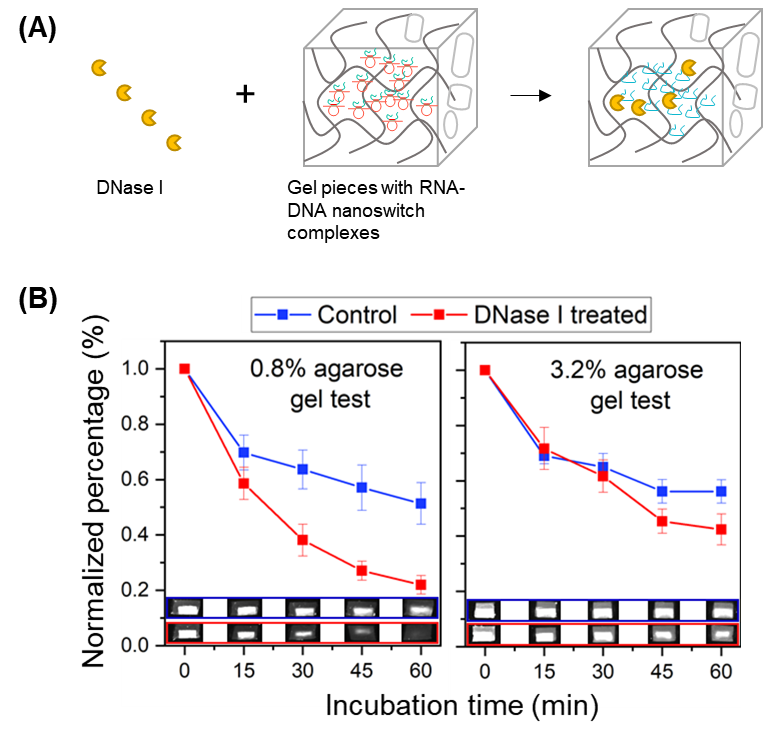

**Figure S1. Digestion of DNA nanoswitches in gel pieces.** (A) Concept of digesting DNA nanoswitches in agarose gel pieces by DNase I. (B) Digestion of DNA nanoswitches in 0.8% (left) and 3.2% (right) agarose gel pieces. 10 ng DNA nanoswitch pre-stained by 1×GelRed (Biotium, Inc.) was loaded to each well. The gel bands of nanoswitches were cut out carefully to ensure similar size and then gel images were taken for reference (shown as inset in the figure). Each gel piece was submerged in 100 µl 1×DNase I reaction buffer (NEB, Inc.) in 1.5 ml tube and then 2U DNase I (NEB, Inc.) was added. Then, all samples were incubated at 37°C. Gel images were taken after incubation for 15, 30, 45 and 60 min.

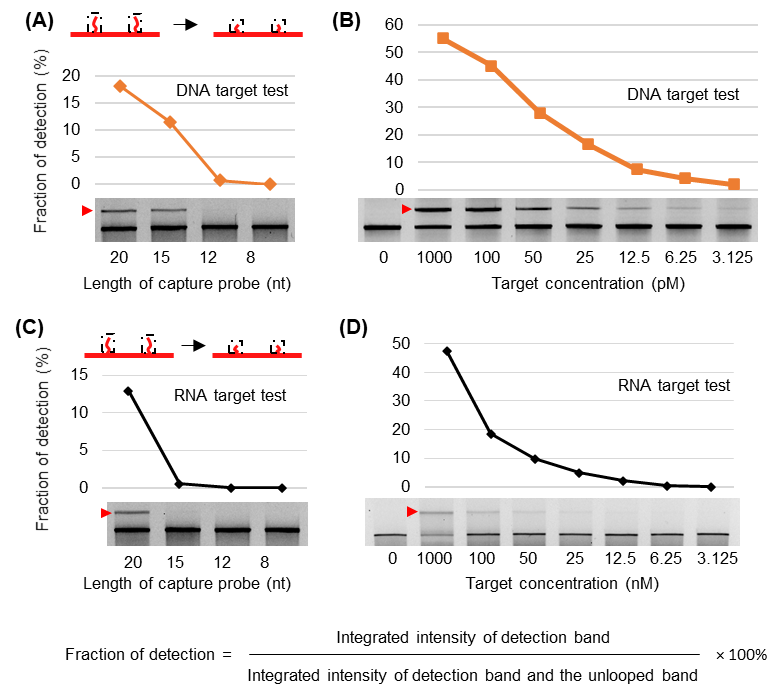

**Figure S2.** **Optimization of the capture probe and sensitivity test of DNA nanoswitch of mRNA fragment.** (**A**) Detection test of the nanoswitch with different capture probe lengths by using corresponding DNA targets shows that the 20 nt probe has the highest detection efficiency. (**B**) Detection sensitivity test of the nanoswitch with 20 nt capture probes by using corresponding DNA targets shows a detection limit as low as 3.1 pM. (**C**) Testing different lengths of capture probes to detect an mRNA fragment. (**D**) Sensitivity of detecting mRNA fragment using a nanoswitch with 20 nt capture probes.

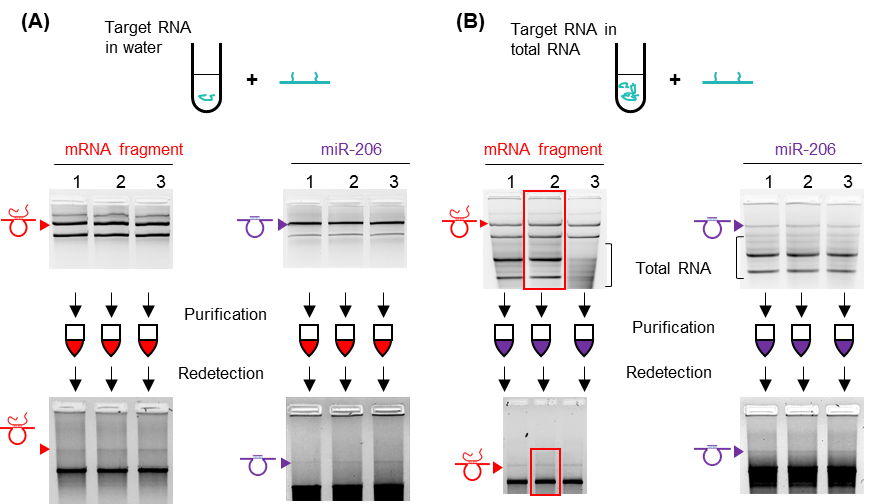

**Figure S3. Proof-of-concept single species RNA purification.** (**A**) Detection, purification and redetection of mRNA fragment and miR-206 in water, finished in triplicate (lanes 1, 2 and 3)**.** (**B**) Detection, purification and redetection of mRNA fragment and miR-206 in total RNA, finished in triplicate (lanes 1, 2 and 3). The red frames indicate the areas of gel images shown in **Figure 1d**. Each pre-purification sample (for each lane) has 30 μl volume that contains 0.53 nM nanoswitch (mRNA fragment) or 0.6 nM nanoswitch (miR-206), 3.3 nM target RNA, 10 mM MgCl_2_, and 1× PBS. 250 ng HeLa total RNA was added for the test within total RNA. Redetection samples contain 10 µL volume with 0.15 nM nanoswitch, 4 µL purified RNA sample, 10 mM MgCl_2_, 1× PBS, and 200 nM blocking oligos.

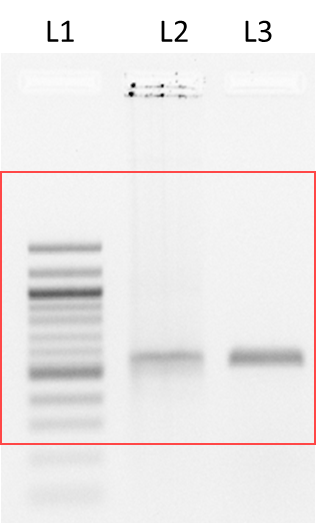

**Figure S4. Length validation for purified mRNA fragment.** The amplicons of qRT-PCR of the purified mRNA fragment is shown in lane L3, alongside the dsDNA template in lane L2 and a 100 bp ladder in lane L1. The gel was 1.6% agarose gel, run in cold room at 60V for 60 min. The frame indicate the areas of gel images shown in **Figure 1d**.

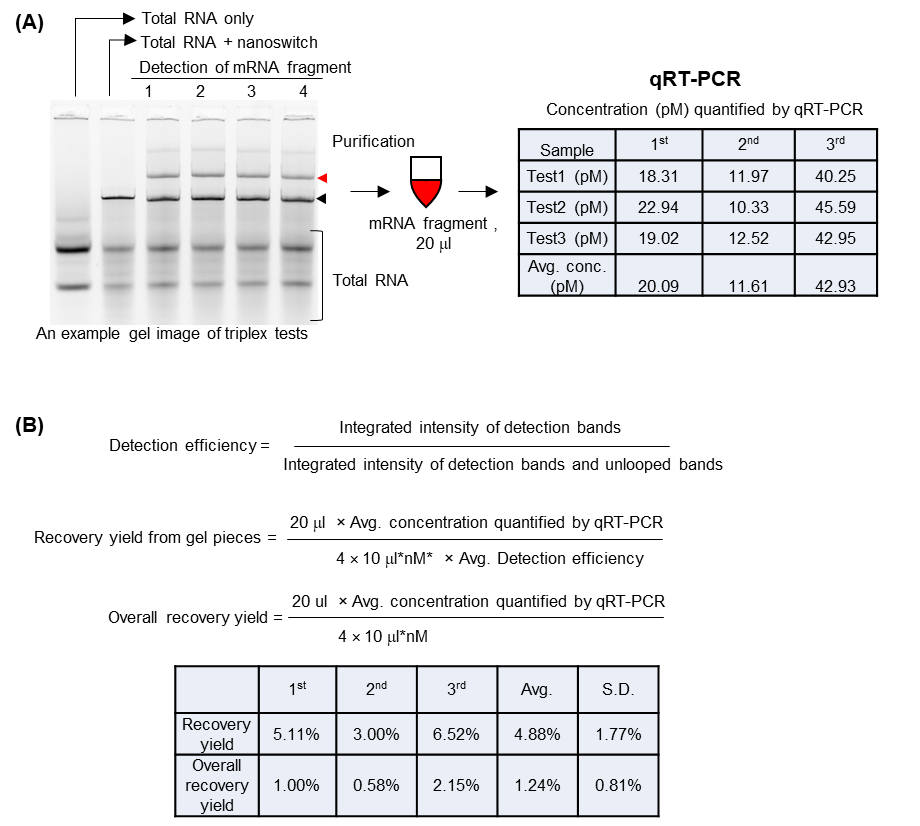

**Figure S5. Quantification of the purification yield of mRNA fragment based on qRT-PCR test.** (A) An example of one of three gel images of mRNA fragment detection. For each test, we combined the detection bands of four lanes (1, 2, 3 and 4), each containing 10 fmol nanoswitches and 10 fmol target mRNA fragment with 10 mM MgCl_2_ and 1× PBS in a 10 µL volume. The results of qRT-PCR of the three independently purified products is shown in the table. Each purified sample was quantified three times by qRT-PCR (test 1 to 3) and the average concentration is the average value of the three tests. (B) Equations used to calculate the recovery yield and overall yield. The data shown in **Figure 1e** is summarized in the table on the bottom.

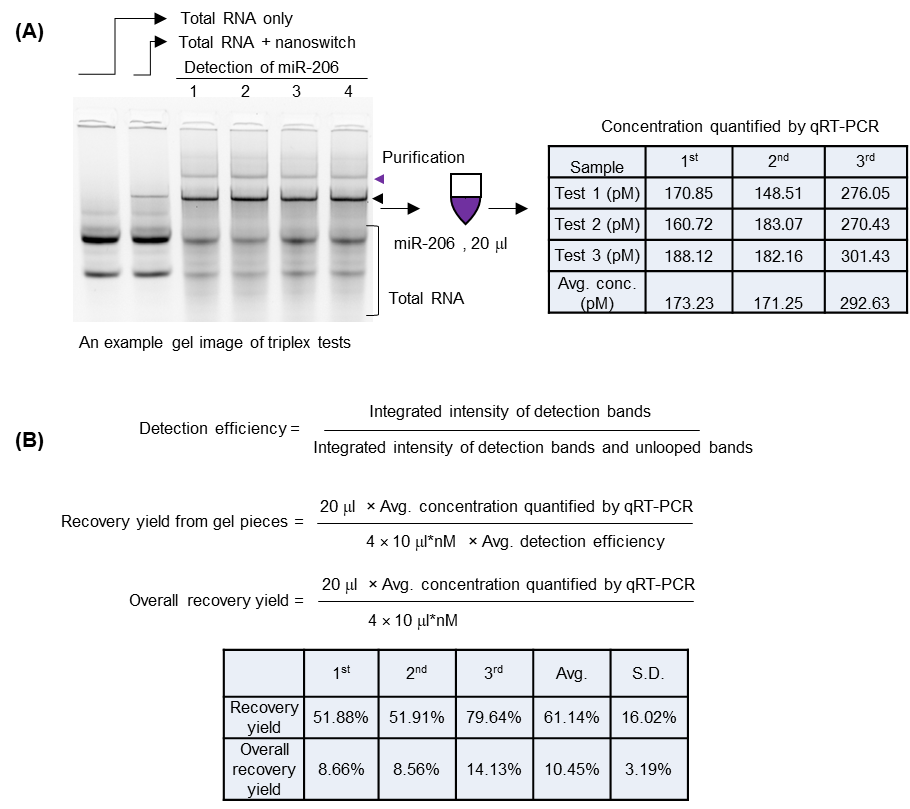

**Figure S6. Quantification of the purification yield of miR-206 based on qRT-PCR test.** (A) An example of one of three gel images of miR-206 detection. For each test, we combined the detection bands of four lanes (1, 2, 3 and 4), each containing 10 fmol nanoswitches and 10 fmol target microRNA with 10 mM MgCl_2_ and 1× PBS in a 10 µL volume. The results of qRT-PCR of the three independently purified products is shown in the table. Each purified sample was quantified three times (test 1 to 3) and average concentration is the average value of the three tests. (B) Equations used to calculate the recovery yield and overall yield. µl*nM: volume × concentration. The data shown in **Figure 1E** is summarized in the table on the bottom.

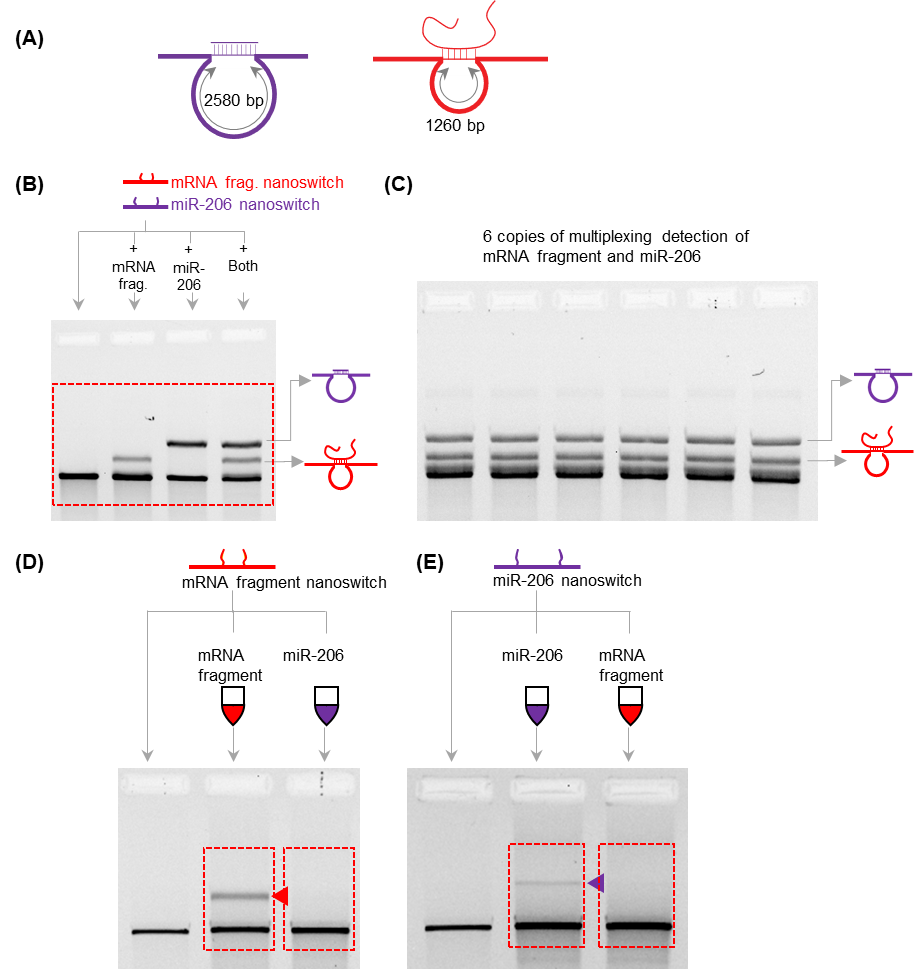

**Figure S7. Gel images of the multiplexed purification of mRNA fragment and miR-206**. (A) The two loop sizes of DNA nanoswitches for mRNA fragment and miR-206 detection. (B) The entire gel image of the multiplexed detection of mRNA fragment and miR-206 in water. The gel was a 0.8% agarose gel and was run in the cold room at 60 V for 2 hours. (C) The entire gel image of the multiplexed detection of mRNA fragment and miR-206 for purification. The gel was a 0.8% agarose gel and was run in the cold room at 65 V for 2 hours. Totally, 60 µl detection sample (containing 0.8 nM unpurified nanoswitch for each target RNA, 10 nM of each target RNA, 10 mM MgCl_2_, 1× PBS) was prepared and 10 µl was loaded to each well. (D-E) Redetection of the mRNA fragment and miR-206 purified from the detection bands shown in (C). The dotted frames indicate the areas of gel images shown in **Figure 1f**.

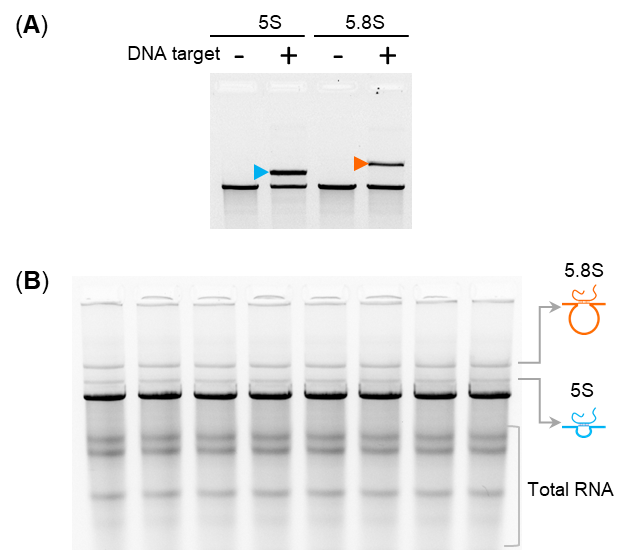

**Figure S8. Detection and purification of 5.8S and 5S rRNA.** (A) Negative control and detection test of nanoswitches designed for 5.8S and 5S rRNA. Detection was validated using DNA controls. (B) The entire gel image of the 5.8S and 5S rRNA multiplexed detection. Totally, 80 µl sample (containining 0.75 nM nanoswitch of each target RNA, 50 ng/µl total RNA of HeLa cell, 10 mM MgCl_2_, 1× PBS, 3.3× GelRed) was prepared and incubated with a thermal annealing ramp (40°C to 25 °C over 12 hours) and 10 µl was loaded to each well. For each target, two clean columns were used and 40 µl purified sample was obtained.

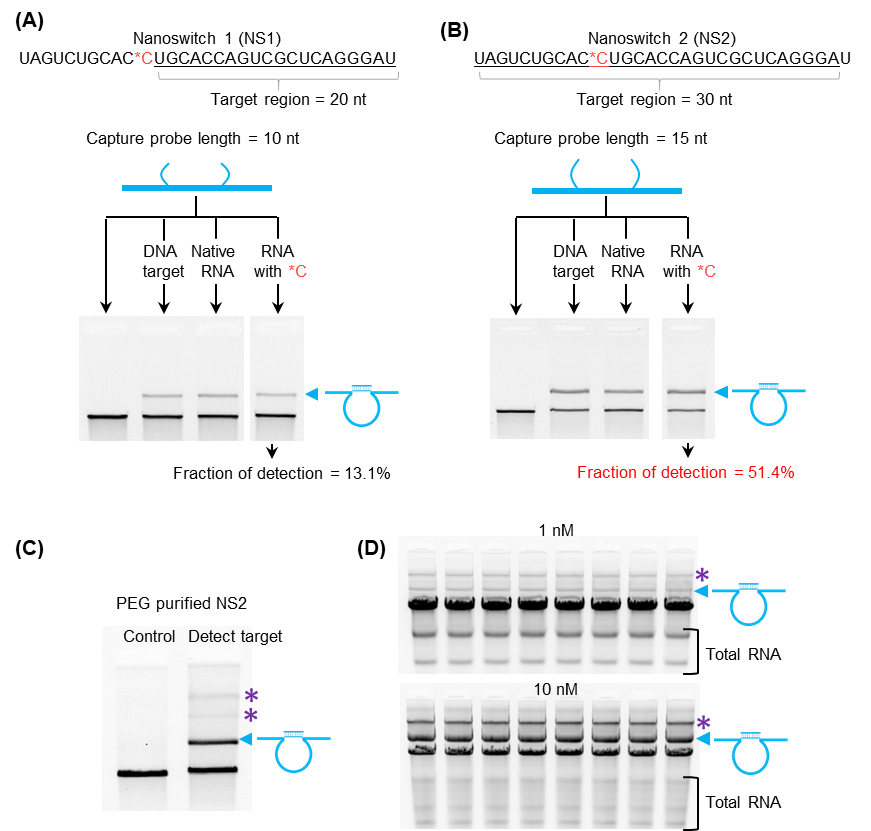

**Figure S9. Purification of RNA with chemical modification.** (A, B) Optimizing design of DNA nanoswitches for the detection of RNA with chemical modification m^4,4^C. The nanoswitch with 15 nt capture probe length has higher detection efficiency compared to the nanoswitch with10 nt capture probes. (C) PEG purification of DNA nanoswitch and the negative control and detection test. (D) Gel images showing detection of chemically-modified RNA (at 1 and 10 nM concentration) when spiked into total RNA. For each case, 80 µl detection sample (containing 4 nM nanoswitch, 50 ng/µl total RNA of HeLa cell, 10 mM MgCl_2_, 1×PBS, 3.3×GelRed) was prepared and incubated with a thermal annealing ramp (40°C to 25 °C over 12 hours) and 10 µl was loaded to each well. * indicates the possible complexes formed by two or more nanoswitches, which is known to occur at high nanoswitch concentration as in this case.

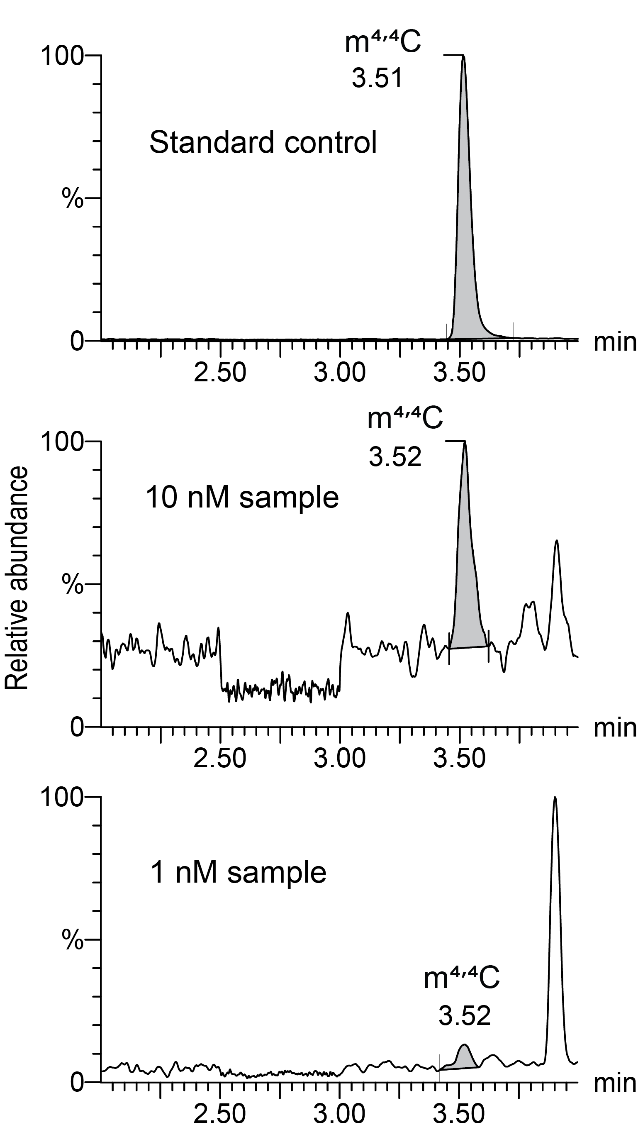

**Figure S10. Purification of RNA molecules with chemical modifications.** Modification probing of the purified RNA molecules by LC-MS/MS, top: m^4,4^C modification standard, middle and bottom are the interrogations of RNA purified from 10 nM and 1 nM samples respectively.

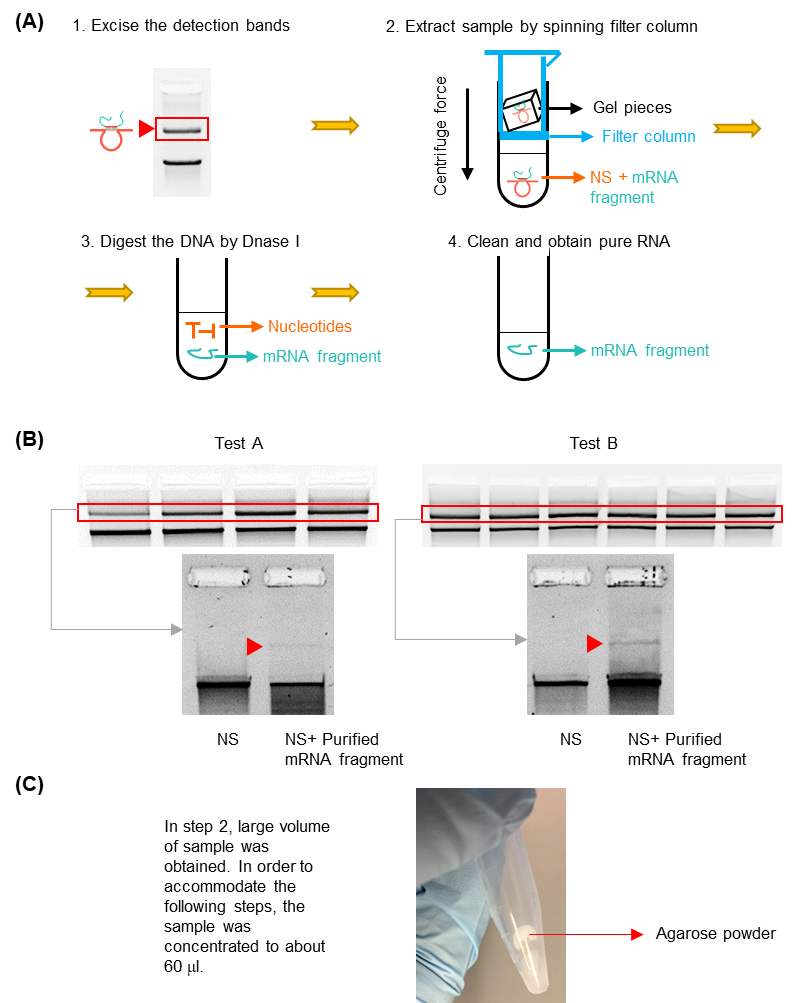

**Figure S11. Test of DNA gel extraction spin columns. (A)** After cutting the gel bands, we used Freeze ‘N Squeeze DNA gel spin columns (Bio-Rad Laboratories, Inc.) to extract RNA-looped DNA nanoswitches for the next purification steps. (**B**) Two tests were conducted and 20 µl purified RNA sample was obtained for each. For verification by redetection, 5 µl purified RNA sample was used. Comparison of the gel band intensities of redetection and initial detection bands shows that the purification yield is low for both tests. (**C**) In step 2, we concentrated the sample extracted from the excised gel pieces in a universal vacuum system (Savant UVS 400) and noticed that after concentration, white powder appeared on the inner wall of the tube. We believe the white powder to be agarose which could influence the following purification steps and result in low purification yield.

**Table S1. Sequence of target mRNA fragment and oligos used for the design of nanoswitches.**

| Name | Sequence (5’->3’) | Len. |
| --- | --- | --- |
| mRNA fragment | CUGGACCUCCCAAAAGCCAACUUAUUGUGAUAUUUGUAAAUUAUAGUUUUAGCAGUUCGUUUGCCACAUGAGUGGAACAUCGUGAAUGCACUUUUGAUAAGUGCUCGGUUAUUUUAUAUUGUAACUACCAGCCUUCAGAGGCGAUCGUAUGCAUAGUUUCUUGAAGUCAAUUUGUCCGUGUAUUCAAAUGUUUGCUUUCGUGAAAACUCGCAUUGUUUUGUCACUCUACCAAGUAAUCAAUUUGUACCAAUCAAUCGCAUAUGGUUGUCCUAGAUCUAAAAAUGGCAAUAAUUUGCCUUCGGUAUUGCACCUAAUGUAUUCAAGAACAAGUAGGG**AAGCUCGAAAUUUCUCAAAUACUUACCCAAAAAAUAGAUA**GAAAUAUAUUUUCGAUUCGCAAUCGU | 401 |
| mRNA frag_DNA Target | AAGCTCGAAATTTCTCAAATACTTACCCAAAAAATAGATA | 40 |
| mRNA_Probe1_20nt | ACCGTTGTAGCAATACTTCTTTGATTAGTAATAACATCACTATCTATTTTTTGGGTAAGT | 60 |
| mRNA_Probe2_20nt_big loop | ATTTGAGAAATTTCGAGCTTTCAACCGATTGAGGGAGGGAAGGTAAATATTGACGGAAAT | 60 |
| mRNA_Probe1_15nt | TGTAGCAATACTTCTTTGATTAGTAATAACATCACATTTTTTGGGTAAGTCAGTC | 55 |
| mRNA_Probe2_15nt_big loop | GTCAGATTTGAGAAATTTCGTCAACCGATTGAGGGAGGGAAGGTAAATATTGACG | 55 |
| mRNA_Probe1_12nt | TGTAGCAATACTTCTTTGATTAGTAATAACATCACTTTTGGGTAAGTCAGTC | 52 |
| mRNA_Probe2_12nt_big loop | GTCAGATTTGAGAAATTTCAACCGATTGAGGGAGGGAAGGTAAATATTGACG | 52 |
| mRNA_Probe1_8nt | TGTAGCAATACTTCTTTGATTAGTAATAACATCACGGGTAAGTCAGTC | 48 |
| mRNA_Probe2_8nt_big loop | GTCAGATTTGAGATCAACCGATTGAGGGAGGGAAGGTAAATATTGACG | 48 |
| mRNA_Probe2_small loop | **ATTTGAGAAATTTCGAGCTT**TGGGTTATATAACTATATGTAAATGCTGATGCAAATCCAA | 60 |

**Table S2. Sequence of target miR-206 and oligos used for the design of nanoswitches.**

| Name | Sequence (5’->3’) | Len. |
| --- | --- | --- |
| miR-206 | UGGAAUGUAAGGAAGUGUGUGG | 22 |
| miR-206 DNA Target | TGGAATGTAAGGAAGTGTGTGG | 22 |
| miR206_Probe1 | ACCGTTGTAGCAATACTTCTTTGATTAGTAATAACATCACCCACACACTTC | 51 |
| miR206_Probe2_big loop | **CTTACATTCCA**TCAACCGATTGAGGGAGGGAAGGTAAATATTGACGGAAAT | 51 |

**Table S3. Sequences of target ribosomal RNA 5.8S and 5S and oligos used for the design of nanoswitches.**

| Name | Sequence (5’->3’) | Len. |
| --- | --- | --- |
| rRNA 5.8S  (NR_146147.1) | CGACUCUUAGCGGUGGAUCACUCGGCUCGUGCGUCGAUGAAGAACGCAGCUAGCUGCGAGAAUUAAUGUGAAUUGCAGGACACAUUGAUCAUCGACACUUCGAACGCACUUGCGGCCCCGGGUUCCUCCCGGGGCUACGCCUGUCUGAGCGUCGCUU | 22 |
| 5.8S_DNA Target | UGGAAUGUAAGGAAGUGUGUGG | 157 |
| 5.8S_Probe1 | ACCGTTGTAGCAATACTTCTTTGATTAGTAATAACATCACTCATCGACGCACGAG | 55 |
| 5.8S_Probe2_big loop | CCGAGTGATCCACCGTCAACCGATTGAGGGAGGGAAGGTAAATATTGACGGAAAT | 55 |
| rRNA 5S  (NR_023376.1) | GUCUACGGCCAUACCACCCUGAACGCGCCCGAUCUCGUCUGAUCUCGGAAGCUAAGCAGGGUCGGGCCUGGUUAGUACUUGGAUGGGAGACCGCCUGGGAAUACCGGGUGCUGUAGGCUUU | 121 |
| 5S_DNA Target | TAGTACTTGGATGGGAGACCGCCTGGGAAT | 30 |
| 5S_Probe1 | ACCGTTGTAGCAATACTTCTTTGATTAGTAATAACATCACATTCCCAGGCGGTCT | 55 |
| 5S_Probe2_big loop | CCCATCCAAGTACTATGGGTTATATAACTATATGTAAATGCTGATGCAAATCCAA | 55 |

**Table S4. Sequence of synthesized RNA with m^4,4^C chemical modification and oligos used for the design of nanoswitches.**

| Name | Sequence (5’->3’) | Len. |
| --- | --- | --- |
| RNA_m^4,4^C | UAGUCUGCACCUGCACCAGUCGCUCAGGGAU | 31 |
| RNA_m^4,4^C_DNA Target | TAGTCTGCACCTGCACCAGTCGCTCAGGGAT | 31 |
| NS1_Probe1 | ACCGTTGTAGCAATACTTCTTTGATTAGTAATAACATCACATCCCTGAGC | 50 |
| NS1_Probe2_big loop | GACTGGTGCATCAACCGATTGAGGGAGGGAAGGTAAATATTGACGGAAAT | 50 |
| NS2_Probe1 | ACCGTTGTAGCAATACTTCTTTGATTAGTAATAACATCACTCCCTGAGCGACTGG | 55 |
| NS2_Probe2_big loop | TGCAGGTGCAGACTATCAACCGATTGAGGGAGGGAAGGTAAATATTGACGGAAAT | 55 |

**Table S5. Backbone used for the design of nanoswitches and blocking oligo used in some of the detection assays**^1,2^**.**

| **Backbone oligos** | | |
| --- | --- | --- |
| **#** | **Sequence (5’-3’)** | **Length** |
| 1 | AGAGCATAAAGCTAAATCGGTTGTACCAAAAACATTATGACCCTGTAATACTTTTGCGGG | 60 |
| 2 | AGAAGCCTTTATTTCAACGCAAGGATAAAAATTTTTAGAACCCTCATATATTTTAAATGC | 60 |
| 3 | AATGCCTGAGTAATGTGTAGGTAAAGATTCAAAAGGGTGAGAAAGGCCGGAGACAGTCAA | 60 |
| 4 | ATCACCATCAATATGATATTCAACCGTTCTAGCTGATAAATTAATGCCGGAGAGGGTAGC | 60 |
| 5 | TATTTTTGAGAGATCTACAAAGGCTATCAGGTCATTGCCTGAGAGTCTGGAGCAAACAAG | 60 |
| 6 | AGAATCGATGAACGGTAATCGTAAAACTAGCATGTCAATCATATGTACCCCGGTTGATAA | 60 |
| 7 | TCAGAAAAGCCCCAAAAACAGGAAGATTGTATAAGCAAATATTTAAATTGTAAACGTTAA | 60 |
| 8 | TATTTTGTTAAAATTCGCATTAAATTTTTGTTAAATCAGCTCATTTTTTAACCAATAGGA | 60 |
| 9 | ACGCCATCAAAAATAATTCGCGTCTGGCCTTCCTGTAGCCAGCTTTCATCAACATTAAAT | 60 |
| 10 | GGATAGGTCACGTTGGTGTAGATGGGCGCATCGTAACCGTGCATCTGCCAGTTTGAGGGG | 60 |
| 11 | ACGACGACAGTATCGGCCTCAGGAAGATCGCACTCCAGCCAGCTTTCCGGCACCGCTTCT | 60 |
| 12 | GGTGCCGGAAACCAGGCAAAGCGCCATTCGCCATTCAGGCTGCGCAACTGTTGGGAAGGG | 60 |
| 13 | CGATCGGTGCGGGCCTCTTCGCTATTACGCCAGCTGGCGAAAGGGGGATGTGCTGCAAGG | 60 |
| 14 | CGATTAAGTTGGGTAACGCCAGGGTTTTCCCAGTCACGACGTTGTAAAACGACGGCCAGT | 60 |
| 15 | GCCAAGCTTGCATGCCTGCAGGTCGACTCTAGAGGATCCCCGGGTACCGAGCTCGAATTC | 60 |
| 16 | GTAATCATGGTCATAGCTGTTTCCTGTGTGAAATTGTTATCCGCTCACAATTCCACACAA | 60 |
| 17 | CATACGAGCCGGAAGCATAAAGTGTAAAGCCTGGGGTGCCTAATGAGTGAGCTAACTCAC | 60 |
| 18 | ATTAATTGCGTTGCGCTCACTGCCCGCTTTCCAGTCGGGAAACCTGTCGTGCCAGCTGCA | 60 |
| 19 | TTAATGAATCGGCCAACGCGCGGGGAGAGGCGGTTTGCGTATTGGGCGCCAGGGTGGTTT | 60 |
| 20 | GTTGCAGCAAGCGGTCCACGCTGGTTTGCCCCAGCAGGCGAAAATCCTGTTTGATGGTGG | 60 |
| 21 | TTCCGAAATCGGCAAAATCCCTTATAAATCAAAAGAATAGCCCGAGATAGGGTTGAGTGT | 60 |
| 22 | TGTTCCAGTTTGGAACAAGAGTCCACTATTAAAGAACGTGGACTCCAACGTCAAAGGGCG | 60 |
| 23 | AAAAACCGTCTATCAGGGCGATGGCCCACTACGTGAACCATCACCCAAATCAAGTTTTTT | 60 |
| 24 | GGGGTCGAGGTGCCGTAAAGCACTAAATCGGAACCCTAAAGGGAGCCCCCGATTTAGAGC | 60 |
| 25 | TTGACGGGGAAAGCCGGCGAACGTGGCGAGAAAGGAAGGGAAGAAAGCGAAAGGAGCGGG | 60 |
| 26 | CGCTAGGGCGCTGGCAAGTGTAGCGGTCACGCTGCGCGTAACCACCACACCCGCCGCGCT | 60 |
| 27 | TAATGCGCCGCTACAGGGCGCGTACTATGGTTGCTTTGACGAGCACGTATAACGTGCTTT | 60 |
| 28 | CCTCGTTAGAATCAGAGCGGGAGCTAAACAGGAGGCCGATTAAAGGGATTTTAGACAGGA | 60 |
| 29 | ACGGTACGCCAGAATCCTGAGAAGTGTTTTTATAATCAGTGAGGCCACCGAGTAAAAGAG | 60 |
| 30 | TTGCCTGAGTAGAAGAACTCAAACTATCGGCCTTGCTGGTAATATCCAGAACAATATTAC | 60 |
| 31 | CGCCAGCCATTGCAACAGGAAAAACGCTCATGGAAATACCTACATTTTGACGCTCAATCG | 60 |
| 32 | TCTGAAATGGATTATTTACATTGGCAGATTCACCAGTCACACGACCAGTAATAAAAGGGA | 60 |
| 33 | CATTCTGGCCAACAGAGATAGAACCCTTCTGACCTGAAAGCGTAAGAATACGTGGCACAG | 60 |
| 34 | ACAATATTTTTGAATGGCTATTAGTCTTTAATGCGCGAACTGATAGCCCTAAAACATCGC | 60 |
| 35 | CATTAAAAATACCGAACGAACCACCAGCAGAAGATAAAACAGAGGTGAGGCGGTCAGTAT | 60 |
| 36 | TAACACCGCCTGCAACAGTGCCACGCTGAGAGCCAGCAGCAAATGAAAAATCTAAAGCAT | 60 |
| 37 | CACCTTGCTGAACCTCAAATATCAAACCCTCAATCAATATCTGGTCAGTTGGCAAATCAA | 60 |
| 38 | CAGTTGAAAGGAATTGAGGAAGGTTATCTAAAATATCTTTAGGAGCACTAACAACTAATA | 60 |
| 39 | GATTAGAGCCGTCAATAGATAATACATTTGAGGATTTAGAAGTATTAGACTTTACAAACA | 60 |
| 40 | CATTATCATTTTGCGGAACAAAGAAACCACCAGAAGGAGCGGAATTATCATCATATTCCT | 60 |
| 41 | GATTATCAGATGATGGCAATTCATCAATATAATCCTGATTGTTTGGATTATACTTCTGAA | 60 |
| 42 | TAATGGAAGGGTTAGAACCTACCATATCAAAATTATTTGCACGTAAAACAGAAATAAAGA | 60 |
| 43 | AATTGCGTAGATTTTCAGGTTTAACGTCAGATGAATATACAGTAACAGTACCTTTTACAT | 60 |
| 44 | CGGGAGAAACAATAACGGATTCGCCTGATTGCTTTGAATACCAAGTTACAAAATCGCGCA | 60 |
| 45 | GAGGCGAATTATTCATTTCAATTACCTGAGCAAAAGAAGATGATGAAACAAACATCAAGA | 60 |
| 46 | AAACAAAATTAATTACATTTAACAATTTCATTTGAATTACCTTTTTTAATGGAAACAGTA | 60 |
| 47 | CATAAATCAATATATGTGAGTGAATAACCTTGCTTCTGTAAATCGTCGCTATTAATTAAT | 60 |
| 48 | TTTCCCTTAGAATCCTTGAAAACATAGCGATAGCTTAGATTAAGACGCTGAGAAGAGTCA | 60 |
| 49 | ATAGTGAATTTATCAAAATCATAGGTCTGAGAGACTACCTTTTTAACCTCCGGCTTAGGT | 60 |
| 50 | GAAAACTTTTTCAAATATATTTTAGTTAATTTCATCTTCTGACCTAAATTTAATGGTTTG | 60 |
| 51 | AAATACCGACCGTGTGATAAATAAGGCGTTAAATAAGAATAAACACCGGAATCATAATTA | 60 |
| 52 | CTAGAAAAAGCCTGTTTAGTATCATATGCGTTATACAAATTCTTACCAGTATAAAGCCAA | 60 |
| 53 | CGCTCAACAGTAGGGCTTAATTGAGAATCGCCATATTTAACAACGCCAACATGTAATTTA | 60 |
| 54 | GGCAGAGGCATTTTCGAGCCAGTAATAAGAGAATATAAAGTACCGACAAAAGGTAAAGTA | 60 |
| 55 | ATTCTGTCCAGACGACGACAATAAACAACATGTTCAGCTAATGCAGAACGCGCCTGTTTA | 60 |
| 56 | TCAACAATAGATAAGTCCTGAACAAGAAAAATAATATCCCATCCTAATTTACGAGCATGT | 60 |
| 57 | AGAAACCAATCAATAATCGGCTGTCTTTCCTTATCATTCCAAGAACGGGTATTAAACCAA | 60 |
| 58 | GTACCGCACTCATCGAGAACAAGCAAGCCGTTTTTATTTTCATCGTAGGAATCATTACCG | 60 |
| 59 | CGCCCAATAGCAAGCAAATCAGATATAGAAGGCTTATCCGGTATTCTAAGAACGCGAGGC | 60 |
| 60 | ATTTTGCACCCAGCTACAATTTTATCCTGAATCTTACCAACGCTAACGAGCGTCTTTCCA | 60 |
| 61 | GAGCCTAATTTGCCAGTTACAAAATAAACAGCCATATTATTTATCCCAATCCAAATAAGA | 60 |
| 62 | AACGATTTTTTGTTTAACGTCAAAAATGAAAATAGCAGCCTTTACAGAGAGAATAACATA | 60 |
| 63 | AAAACAGGGAAGCGCATTAGACGGGAGAATTAACTGAACACCCTGAACAAAGTCAGAGGG | 60 |
| 64 | TAATTGAGCGCTAATATCAGAGAGATAACCCACAAGAATTGAGTTAAGCCCAATAATAAG | 60 |
| 65 | AGCAAGAAACAATGAAATAGCAATAGCTATCTTACCGAAGCCCTTTTTAAGAAAAGTAAG | 60 |
| 66 | CAGATAGCCGAACAAAGTTACCAGAAGGAAACCGAGGAAACGCAATAATAACGGAATACC | 60 |
| 67 | CAAAAGAACTGGCATGATTAAGACTCCTTATTACGCAGTATGTTAGCAAACGTAGAAAAT | 60 |
| 68 | ACATACATAAAGGTGGCAACATATAAAAGAAACGCAAAGACACCACGGAATAAGTTTATT | 60 |
| 69 | TTGTCACAATCAATAGAAAATTCATATGGTTTACCAGCGCCAAAGACAAAAGGGCGACAT | 60 |
| 70 | TCACCGTCACCGACTTGAGCCATTTGGGAATTAGAGCCAGCAAAATCACCAGTAGCACCA | 60 |
| 71 | TTACCATTAGCAAGGCCGGAAACGTCACCAATGAAACCATCGATAGCAGCACCGTAATCA | 60 |
| 72 | GTAGCGACAGAATCAAGTTTGCCTTTAGCGTCAGACTGTAGCGCGTTTTCATCGGCATTT | 60 |
| 73 | TCGGTCATAGCCCCCTTATTAGCGTTTGCCATCTTTTCATAATCAAAATCACCGGAACCA | 60 |
| 74 | GAGCCACCACCGGAACCGCCTCCCTCAGAGCCGCCACCCTCAGAACCGCCACCCTCAGAG | 60 |
| 75 | CCACCACCCTCAGAGCCGCCACCAGAACCACCACCAGAGCCGCCGCCAGCATTGACAGGA | 60 |
| 76 | GGTTGAGGCAGGTCAGACGATTGGCCTTGATATTCACAAACAAATAAATCCTCATTAAAG | 60 |
| 77 | CCAGAATGGAAAGCGCAGTCTCTGAATTTACCGTTCCAGTAAGCGTCATACATGGCTTTT | 60 |
| 78 | GATGATACAGGAGTGTACTGGTAATAAGTTTTAACGGGGTCAGTGCCTTGAGTAACAGTG | 60 |
| 79 | CCCGTATAAACAGTTAATGCCCCCTGCCTATTTCGGAACCTATTATTCTGAAACATGAAA | 60 |
| 80 | CCAGGCGGATAAGTGCCGTCGAGAGGGTTGATATAAGTATAGCCCGGAATAGGTGTATCA | 60 |
| 81 | CCGTACTCAGGAGGTTTAGTACCGCCACCCTCAGAACCGCCACCCTCAGAACCGCCACCC | 60 |
| 82 | TCAGAGCCACCACCCTCATTTTCAGGGATAGCAAGCCCAATAGGAACCCATGTACCGTAA | 60 |
| 83 | CACTGAGTTTCGTCACCAGTACAAACTACAACGCCTGTAGCATTCCACAGACAGCCCTCA | 60 |
| 84 | TAGTTAGCGTAACGATCTAAAGTTTTGTCGTCTTTCCAGACGTTAGTAAATGAATTTTCT | 60 |
| 85 | GTATGGGATTTTGCTAAACAACTTTCAACAGTTTCAGCGGAGTGAGAATAGAAAGGAACA | 60 |
| 86 | ACTAAAGGAATTGCGAATAATAATTTTTTCACGTTGAAAATCTCCAAAAAAAAGGCTCCA | 60 |
| 87 | AAAGGAGCCTTTAATTGTATCGGTTTATCAGCTTGCTTTCGAGGTGAATTTCTTAAACAG | 60 |
| 88 | CTTGATACCGATAGTTGCGCCGACAATGACAACAACCATCGCCCACGCATAACCGATATA | 60 |
| 89 | TTCGGTCGCTGAGGCTTGCAGGGAGTTAAAGGCCGCTTTTGCGGGATCGTCACCCTCAGC | 60 |
| 90 | CTTTTTCATGAGGAAGTTTCCATTAAACGGGTAAAATACGTAATGCCACTACGAAGGCAC | 60 |
| 91 | CAACCTAAAACGAAAGAGGCAAAAGAATACACTAAAACACTCATCTTTGACCCCCAGCGA | 60 |
| 92 | TTATACCAAGCGCGAAACAAAGTACAACGGAGATTTGTATCATCGCCTGATAAATTGTGT | 60 |
| 93 | CGAAATCCGCGACCTGCTCCATGTTACTTAGCCGGAACGAGGCGCAGACGGTCAATCATA | 60 |
| 94 | AGGGAACCGAACTGACCAACTTTGAAAGAGGACAGATGAACGGTGTACAGACCAGGCGCA | 60 |
| 95 | TAGGCTGGCTGACCTTCATCAAGAGTAATCTTGACAAGAACCGGATATTCATTACCCAAA | 60 |
| 96 | TCAACGTAACAAAGCTGCTCATTCAGTGAATAAGGCTTGCCCTGACGAGAAACACCAGAA | 60 |
| 97 | CGAGTAGTAAATTGGGCTTGAGATGGTTTAATTTCAACTTTAATCATTGTGAATTACCTT | 60 |
| 98 | ATGCGATTTTAAGAACTGGCTCATTATACCAGTCAGGACGTTGGGAAGAAAAATCTACGT | 60 |
| 99 | TAATAAAACGAACTAACGGAACAACATTATTACAGGTAGAAAGATTCATCAGTTGAGATT | 60 |
| 100 | TAAGAGCAACACTATCATAACCCTCGTTTACCAGACGACGATAAAAACCAAAATAGCGAG | 60 |
| 101 | AGGCTTTTGCAAAAGAAGTTTTGCCAGAGGGGGTAATAGTAAAATGTTTAGACTGGATAG | 60 |
| 102 | CGTCCAATACTGCGGAATCGTCATAAATATTCATTGAATCCCCCTCAAATGCTTTAAACA | 60 |
| 103 | GTTCAGAAAACGAGAATGACCATAAATCAAAAATCAGGTCTTTACCCTGACTATTATAGT | 60 |
| 104 | CAGAAGCAAAGCGGATTGCATCAAAAAGATTAAGAGGAAGCCCGAAAGACTTCAAATATC | 60 |
| 105 | GCGTTTTAATTCGAGCTTCAAAGCGAACCAGACCGGAAGCAAACTCCAACAGGTCAGGAT | 60 |
| 106 | TAGAGAGTACCTTTAATTGCTCCTTTTGATAAGAGGTCATTTTTGCGGATGGCTTAGAGC | 60 |
| 107 | TTAATTGCTGAATATAATGCTGTAGCTCAACATGTTTTAAATATGCAACTAAAGTACGGT | 60 |
| 108 | GTCTGGAAGTTTCATTCCATATAACAGTTGATTCCCAATTCTGCGAACGAGTAGATTTAG | 60 |
| 109 | TTTGACCATTAGATACATTTCGCAAATGGTCAATAACCTGTTTAGCTAT | 49 |

| **Variable oligos** | | |
| --- | --- | --- |
| **Name** | **Sequence (5’-3’)** | **Length** |
| Var 1 | AACATCCAATAAATCATACAGGCAAGGCAAAGAATTAGCAAAATTAAGCAATAAAGCCTC | 60 |
| Var 2 | GTGAGCGAGTAACAACCCGTCGGATTCTCCGTGGGAACAAACGGCGGATTGACCGTAATG | 60 |
| Var 3 | TTCTTTTCACCAGTGAGACGGGCAACAGCTGATTGCCCTTCACCGCCTGGCCCTGAGAGA | 60 |
| **Var 4** | TCTGTCCATCACGCAAATTA**ACCGTTGTAGCAATACTTCTTTGATTAGTAATAACATCAC** | **60** |
| Var 5 | **ATTCGACAACTCGTATTAAATCCTTTGCCCGAACGTTATT**AATTTTAAAAGTTTGAGTAA | 60 |
| **Var 6** | **TGGGTTATATAACTATATGTAAATGCTGATGCAAATCCAA**TCGCAAGACAAAGAACGCGA | 60 |
| **Var 7** | **GTTTTAGCGAACCTCCCGACTTGCGGGAGGTTTTGAAGCC**TTAAATCAAGATTAGTTGCT | 60 |
| **Var 8** | **TCAACCGATTGAGGGAGGGAAGGTAAATATTGACGGAAAT**TATTCATTAAAGGTGAATTA | **60** |
| Var 9 | GTATTAAGAGGCTGAGACTCCTCAAGAGAAGGATTAGGATTAGCGGGGTTTTGCTCAGTA | 60 |
| Var 10 | AGCGAAAGACAGCATCGGAACGAGGGTAGCAACGGCTACAGAGGCTTTGAGGACTAAAGA | 60 |
| Var 11 | TAGGAATACCACATTCAACTAATGCAGATACATAACGCCAAAAGGAATTACGAGGCATAG | 60 |
| Var 12 | ATTTTCATTTGGGGCGCGAGCTGAAAAGGTGGCATCAATTCTACTAATAGTAGTAGCATT | 60 |

| **Filler oligos** | | |
| --- | --- | --- |
| **Name** | **Sequence (5’-3’)** | **Length** |
| Var 4 filler | TCTGTCCATCACGCAAATTA | 20 |
| Var 5 filler | AATTTTAAAAGTTTGAGTAA | 20 |
| Var 6 filler | TCGCAAGACAAAGAACGCGA | 20 |
| Var 7 filler | TCGCAAGACAAAGAACGCGA | 20 |
| Var 8 filler | TATTCATTAAAGGTGAATTA | 20 |
| Var 9 filler | TAGCGGGGTTTTGCTCAGTA | 20 |

| **Other oligos** | |  |
| --- | --- | --- |
| Blocking | TCTCATGGCCCTTC | 14 |
| BtsCI cut site oligo | CTACTAATAGTAGTAGCATTAACATCCAATAAATCATACA | 40 |

**Table S6. Cost of the DNA nanoswitch assay**^1,2^**.**

| **Material** | **Unit cost** | **Amount needed/lane** | **Cost per reaction** | **Vendor** |
| --- | --- | --- | --- | --- |
| ***DNA*** | | | | |
| Scaffold DNA | $ 3.40 /µg | 0.0024 µg | $ 0.008 | NEB |
| Oligonucleotides | $ 0.37 /µg | 0.0024 µg | $ 0.001 | IDT |
| Linearization enzyme | $ 0.68 /µL | 0.0032 µL | $ 0.002 | NEB |
| ***Gel reagents*** | | | | |
| Agarose | $ 0.84 /g | 0.02 g | $ 0.01680 | Sigma |
| GelRed | $ 0.23 /µL | 0.0001 µL | $ 0.00002 | Biotium |
| Ficoll (loading dye) | $ 0.21 /g | 0.0003 g | $ 0.00006 | VWR |
| ***Total*** |  |  | $ 0.02788 |  |
